# Human force control may trade-off force error with central tendency and recency biases

**DOI:** 10.1101/2023.09.19.558388

**Authors:** Hansol X. Ryu, Manoj Srinivasan

## Abstract

Understanding how humans control force is useful for understanding human movement behaviors and sensorimotor control. However, it is not well understood how the human nervous system handles different control criteria such as accuracy and energetic cost. We conducted force tracking experiments where participants applied force isometrically while receiving visual force feedback, tracking step changes in target forces. The experiments were designed to disambiguate different plausible objective function components. We found that force tracking error was largely explained by a trade-off between error-reducing tendency and force biases, but we did not need to include an effort-saving tendency. Central tendency bias, which is a shift towards the center of the task distribution, and recency bias, which is a shift towards recent action, were necessary to explain many of our observations. Surprisingly, we did not observe such biases when we removed force requirements for pointing to the target, suggesting that such biases may be task-specific. This study provides insights into the broader field of motor control and human perceptions where behavioral or perceptual biases are involved.

---

We interact with the world by applying forces. For example, we push the ground to walk, press pedals to ride a bicycle, and apply muscle forces onto our own body parts to breathe, sing, and reach out a hand. Understanding how people control forces would be beneficial in understanding how the nervous system performs sensorimotor control, and insightful in designing robots and biomechanical simulations. One way to model sensorimotor control is to view it as an optimization problem, that there are objective functions that are optimized for and constraints that are needed to be satisfied while executing the task (***Baron and Kleinman, 1969***; ***Kleinman et al., 1970***). There often are multiple objectives and constraints in many sensorimotor tasks and robotics applications (***Dao et al., 2016***; ***Jin et al., 2021***). Our goal in this research was to observe human subjects’ behaviors when they perform simple force tracking tasks, and to investigate objective functions that could explain the observed behaviors.

In biomechanics simulations and robotics, researchers often formulate an objective function as a sum of error (or performance) term and effort (or energy cost, control energy) term. Such formulations are used to model human motor control (***Emken et al., 2007***; ***Izawa et al., 2008***; ***Mi et al., 2009***), in biomechanics simulations (***De Groote et al., 2016***; ***Lee and Umberger, 2016***), and in robotics (***Kalakrishnan et al., 2013***; ***Miao et al., 2021***). In optimal control theory, cost function of a linear quadratic regulator (“LQR”, ***Kalman et al. (1960)***) is often formulated as a sum of quadratic functions of state and control input.

Having an error term in the objective function ensures that the goal is achieved to a certain degree. In addition to the error term, there are several reasons to include an effort term, including: 1) To handle the redundancy problem: there are usually redundancies in the system, meaning there are multiple ways to achieve the goal, so additional criteria are needed (as discussed in ***De Groote and Falisse (2021)***). 2) To better mimic biological systems: there are pieces of evidence suggesting that biological systems minimize energetic cost, so having the term allows simulation and robot to behave more like biological systems and thus provides more insights into understanding them (e.g., ***Srinivasan (2011)***). 3) For practical reasons: power or energy consumption are often some of the major concerns for robot designs (e.g., ***Liu and Sun (2013)***; ***Pellegrinelli et al. (2015)***). However, those functions and their relative weightings are usually arbitrarily designed and tuned until they produce acceptable outcomes. Some researchers refer to biological measurements to formulate some components of the objective function (e.g., ***Körding and Wolpert (2004)*** studied error, ***Berret et al. (2011)*** studied cost), but such investigations have not been considered in the context of human force tracking.

We designed a force tracking experiment that allows us to compare various objective function models to experimental observations. Participants were asked to apply forces isometrically onto the platform in front of them through their hands, while receiving the force feedback as the height of the bar on the screen (figure 1A, B). The force targets (figure 1C) they tracked were either clearly defined as a fixed single dot, or vaguely defined in two ways, as a single dot that had noise in its location or as a cloud of multiple dots. The targets were shown for 4 seconds (referred to as “sub-trial”), and the next target of the same kind appeared at a random location. We analyzed force error near the end of each sub-trials (figure 1D) to quantify the force tracking error.

**Figure 1.**
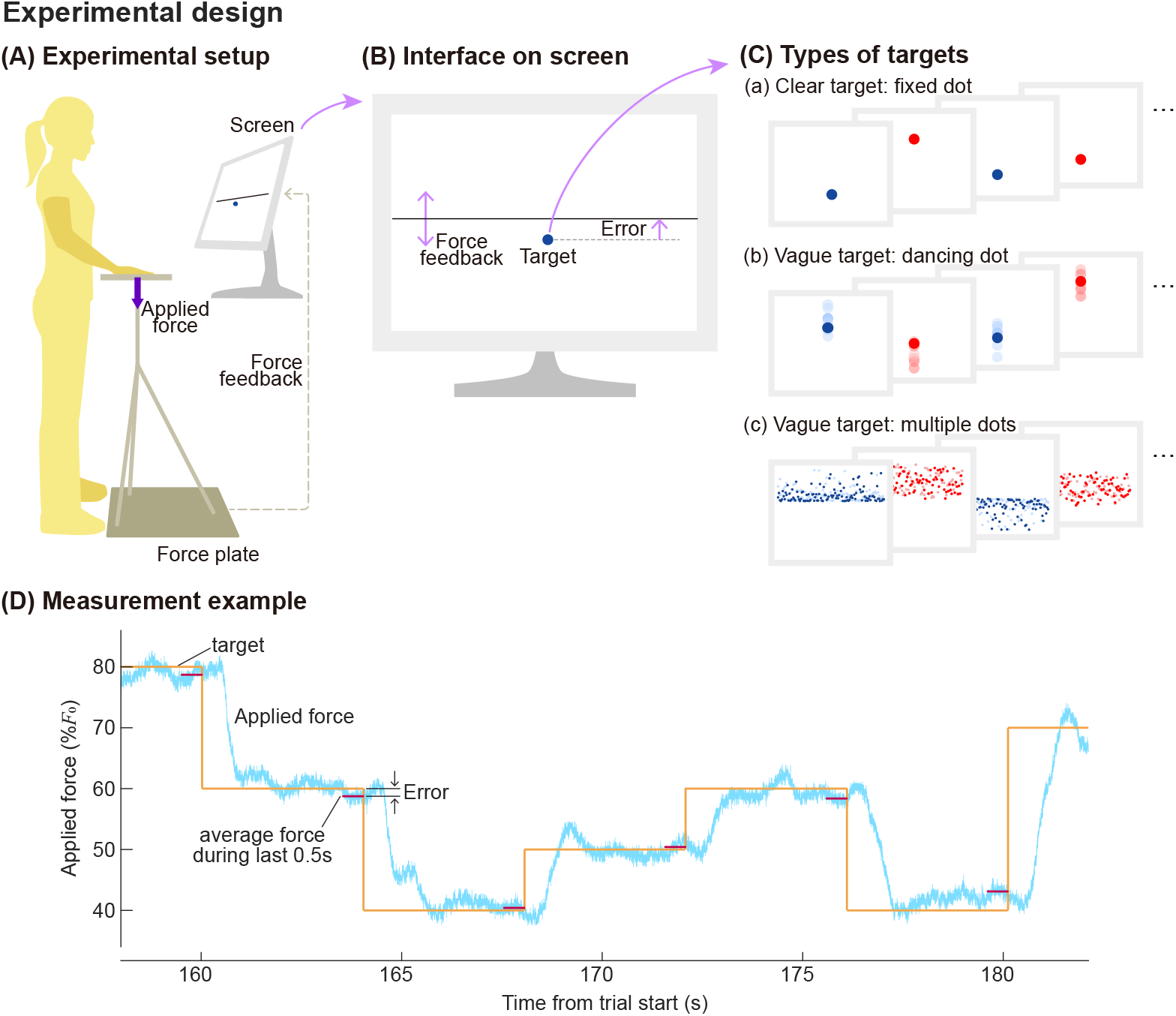
Experimental design. (A) Subjects applied force on the platform in front of them while looking at a screen. The vertical force they applied was measured through a ground-mounted force plate, and the force was relayed to a (B) screen that displayed the applied force as a horizontal bar of a changing vertical location, as well as a target. (C) There were three types of targets. Illustrated are conceptual representations of each sub-trials. (D) An example of a trial, which was a series of sub-trials that changed target force in a random order. Applied force and target force at the time are shown together. Force tracking error was defined as a difference between the average applied force during the last 0.5 seconds of the sub-trial and the mode of the target.

We hypothesized that 1) there would be increasingly negative force tracking errors when the force requirement increases, and 2) the vagueness of the target would result in more shifts in subjects’ behaviors. Consider the case when subjects see the same distribution of dots simply shifted up to require more force (figure 2A). If unlike our hypothesis, effort-saving is not a control criterion, and error-reduction largely explains the behaviors, subjects would increase the force by the same amount as the target shift, and force tracking error would be independent of the target force level (figure 2B-1). However, if they save effort, subjects will make less shift than the target shift, thus making increasingly negative errors as the target force level increases (figure 2B-2). In that case, the behavior can be modeled as a trade-off between error-reducing and effort-saving criteria in the objective function. Additionally, we speculated that the vagueness of the target could stimulate more behavioral changes towards the error-saving side.

**Figure 2.**
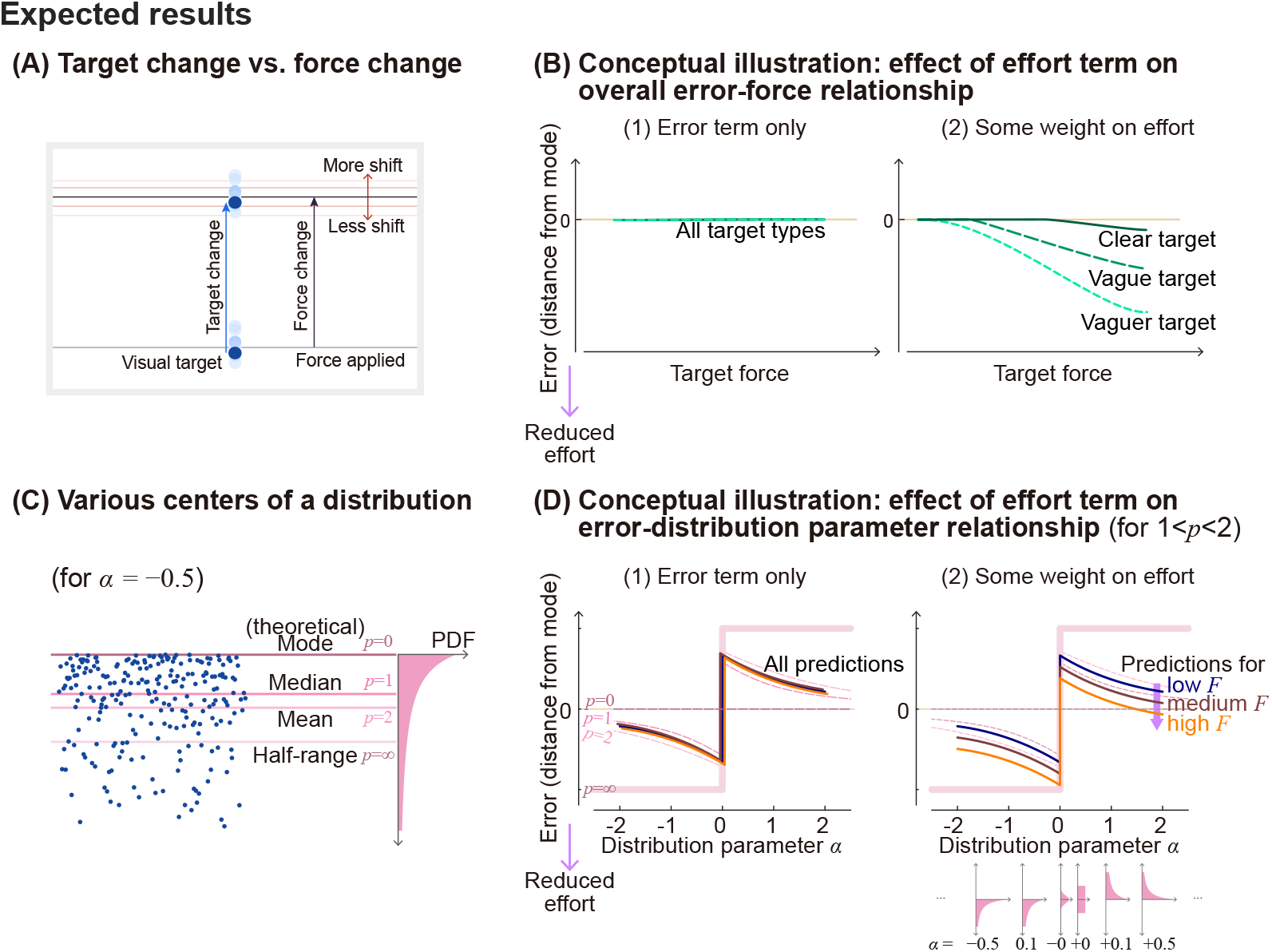
Illustration of expected results for different target types and distributions.(A) Relative force shifts with respect to the target shift. Subjects may shift by the same degree as the visual target position changes, but they may shift less than the ascending target change if effort becomes a critical factor. (B) Therefore, if minimizing error is the only factor that determines people’s behaviors, (1) we expect an average zero error for all force ranges and target types. However, if effort is a consideration too, and if vagueness of the target is an extra allowance to facilitate effort-saving behavior, (2) we expect increasing error towards the effort-saving direction as target vagueness increases. (C) Using targets of different distributions could provide us with extra information about how people perceived the task. As there are many definitions of centers in a skewed distribution, including the mode, median, mean, and half-range, we could study where people were aiming for, and whether this changes with increasing force demands. (D) Combined with effort considerations, if effort does not affect the behavior and people are consistent with what “center” they aim for, (1) we expect to see an error trend across various distribution parameters that is best described by which center they aimed for, and does not change with increased force requirements. If effort is combined with this perception, (2) we expect that people’s response will be similar to that of (1) when force requirement is low, but will shift towards effort-saving way as force requirement increases.

In addition, we used skewed distributions when we defined vague targets to study the error term of the objective function (figure 2C). Skewed distributions have distinct mode, median, mean, and half of the range, which are the locations where error raised to power of 0, 1, 2, and infinity are minimized. For example, if a subject minimized the error raised to about 1.5, on average they would place the force feedback bar between the median and mean of the distributions (figure 2D-1). If an effort-saving tendency is added to the same error-reducing criterion in the objective function, we expect that people’s behaviors would not change much in a low force range, but it will shift towards a direction that saves effort when the force requirement increases (figure 2D-2). We measured how participants changed their force according to the force requirement and given distributions, and applied inverse-optimization analysis to look for an objective function that best describes observed human behaviors.

In sum, we observed increasingly negative force tracking errors in the medium to high force range. However, there also were unexpected findings that looked like people systematically “wasted” effort in the low to medium force range, meaning, they exerted more force than needed for no obvious benefits in reducing errors. We used “central tendency bias” and “recency bias” to explain the results, and conducted additional follow-up experiments on subsets of the participants to test these biases. We use the term “central tendency bias” to describe the shift towards the center of the tasks. That is, if their current force is on a lower side of the experimental range, people tend to make errors towards the center and end up producing more force than needed. In a similar manner, we use the term “recency bias” to refer to the bias towards the recent past action, meaning people tend to make an incomplete shift to the next target. Surprisingly, an effort term that increases with force, does not seem to have a significant effect on the model prediction. Further, we found that eliminating the force requirement as much as possible eliminated the biases, showing no significant bias in visual perception of the tasks.

## Results

### How humans track step changes in forces

Force tracking errors have task-dependent positive and negative biases Subjects typically took about 0.4 s seconds to initiate the force change after the target location changed, and held the force until the next target was shown (figure 1D; average responses can be found in figure 9 and figure 10). We report the difference between the applied force and mode of the target distribution as the tracking error in this paper unless otherwise noted.

We measured the force tracking error for different target types and force requirements. As we expected, people exerted less force relative to the target when the target force increased, and there was a bigger change when the target was vaguer (figure 3A). However, what was unexpected for us was that people tended to exert more force than needed for the low-medium target force range. These two observations, 1) negative correlation between error and target force and 2) positive error on the lower target force, were consistent across subjects (figure 3B). When targets were grouped based on the distribution parameters, force errors were in general in between the median and half-range of the distributions. Each of the sub-grouped forces errors showed a negative correlation with a target force (figure 3C).

**Figure 3.**
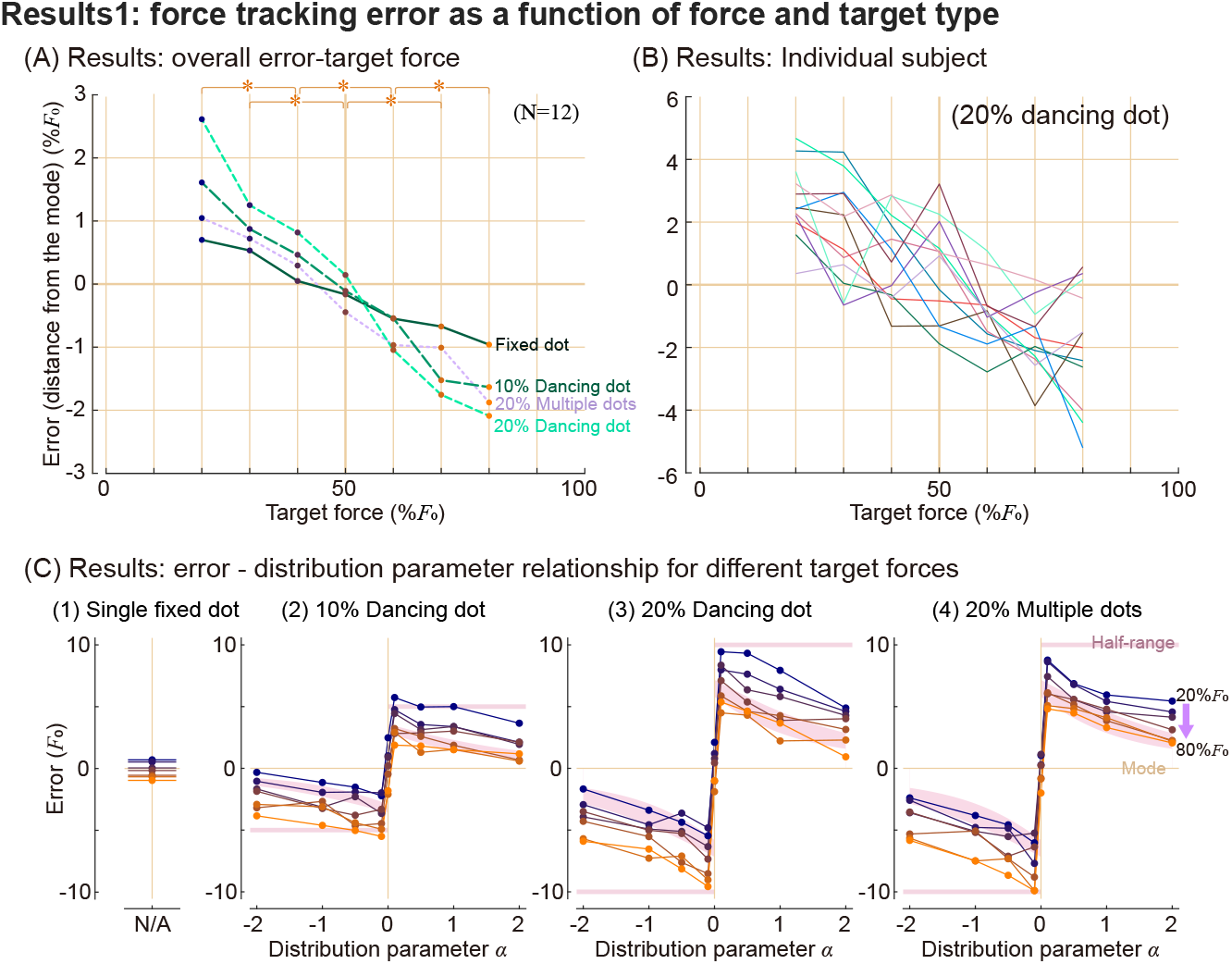
Median values of tracking error for different target force, target types, and distribution parameters. (A) Error, defined as a distance between the average force people applied during the last 0.5 seconds of sub-trials and a mode of the target, for each target type. Asterisks indicate statistically significant differences (p<0.05) for pairs of the force conditions, for every target type compared separately. (B) Error-force pattern of individual subjects. (C) Median response for each target force and distribution parameter. Pink shaded regions represent the range between the theoretical mean and median of each distribution, and pink horizontal lines represent half of the distribution range.

### Force tracking error results are not explained by error and effort minimization

We used two types of objective function models to describe the observations. The objective function we originally formulated was in the form of ∑error^*p*^/*N* + *cF*^Y^, where error is the absolute distance between the target and applied force, *F* is applied force, *N* is the number of target dots. Hyperparameters *p* and *γ* are shape parameters for each functions, and *c* is a constant that determines the relative weighting between two terms. Since human data is noisy, we aimed to keep the formulation as simple as possible. For simplicity, we used ∑(error^*p*^)/*N* to represent performance criteria for targets of different vagueness. The formulation has shallower minima for bigger *N* if *p* < 2, thus competing terms in the objective function would make the minima shift more for bigger *N*. The effort term in this form has inherent limitations, that it cannot explain positive force errors (figure 4A), and thus inverse optimization did not have local minima within our search range. We illustrated a solution that was found on one end of the search range, but even bigger exponents on effort term would further minimize the objective function. Bigger exponents would make low to middle-range errors closer to zero, thus reducing model error. However, this formulation does not predict positive error in any case.

**Figure 4.**
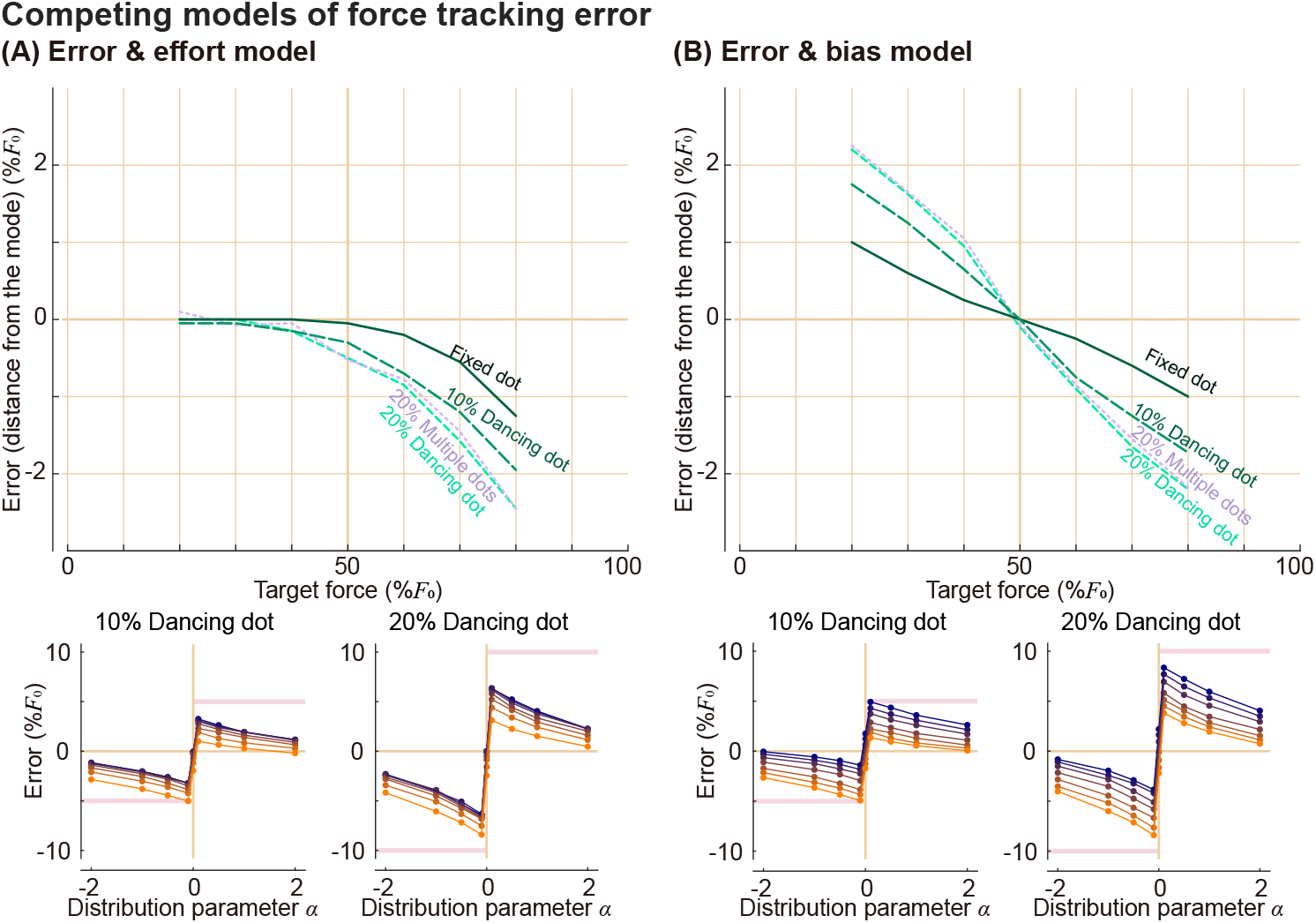
Modeling force tracking errors using (A) error-and-effort model and (B) error-and-bias model. In error-force plots (top row), different target types are indicated with different line types. In distribution parameter-error plots (bottom row), lighter and more orange colors indicate lower target forces, darker and more blue colors indicate higher target forces. Errors shown on the y-axis in both types of plots are both errors with respect to the mode of the target distribution. Objective function models to generate the model prediction here are ∑error^1.6^/*N* + 0.065*F*^5^ for (A) and 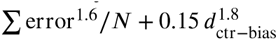 for (B).

### We investigated which variable predicts force tracking error

Force tracking error might be fully determined by the target force or visual representation of the target force. However, there might be a bias that is determined by the temporal relationships between targets. We formulated variants of the objective function models to represent these cases. One of our objective function models was error-and-bias model (figure 4B), which had a form of 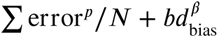, where *d*_bias_ represents the distance to some point when there is a tendency to shift towards that point, which will be further explained later. error is absolute distance between the target and applied force. Hyperparameters *p* and *β* are shape parameters for each functions, and *b* is a constant that determines the relative weighting between two terms. figure 4B shows the error-and-bias model that has a shift towards the average target force.

### Additional experiments to disambiguate different terms

Some of the variables inherently co-varied in the original protocol (referred to as “Protocol 1”), so we designed additional experiments to distinguish some of them and test models accordingly.

The design of the additional protocol and what we found from each of them are listed below. A subset of subjects who participated in the original experiment using Protocol 1 participated in these additional tests, so we compared their responses to those of Protocol 1. We used a single fixed dot as a target in all additional tests to reduce the variability due to the target vagueness.

### Protocol 1 – reduced *F* shows that target force alone does not determine error

In Protocol 1 – reduced *F*, we tested the lower half range of the force while keeping the range of the visual target fixed (figure 5A). Among our 12 subjects who participated in the Protocol 1 experiment, 6 of them participated in this additional protocol experiment on the same day.

**Figure 5.**
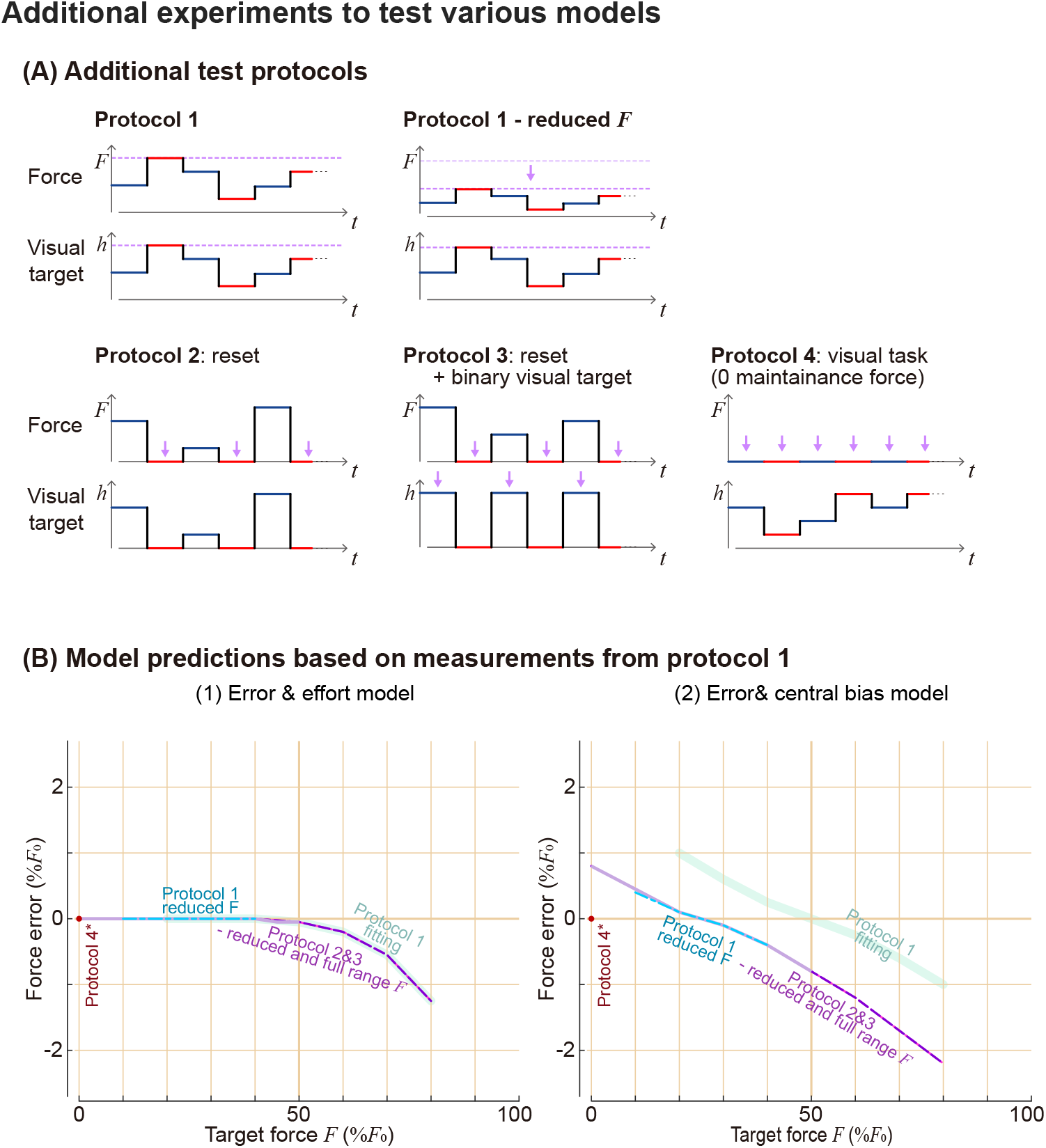
Additional protocols to test different model predictions. (A) Test protocols in terms of force requirement and its visual representation on the screen. Purple arrows emphasize the main differences between protocols. Protocol 1-reduced *F* tested for a lower half range of the force, while spanning to a full visual target range. Protocols 2 and 3 had zero re-set period between each non-zero target force. Protocol 3 had binary targets, where the target was alternating between only two locations each indicating zero and non-zero target. This was done by changing the conversion ratio between force and visual target location each time. Protocol 4 was done using a computer mouse, and did not require force to maintain the pointing location. (B) Example of model predictions for these additional protocols. (1) Error-and-effort model, as an example of a model where target forces determine the errors: results from all protocols will match with each other when the target force is the same. (2) Error-and-bias model, as an example of a model that depends on a target force and the experimental contexts. Error-force trend changes for each protocol, and we had specific qualitative expectations for each protocol.

## Results

Testing result shows that force-error relationship was significantly different between Protocol 1 and Protocol 1 – reduced *F*, but visual target location – error relationship in general matched quite well between the two protocols (figure 6A).

**Figure 6.**
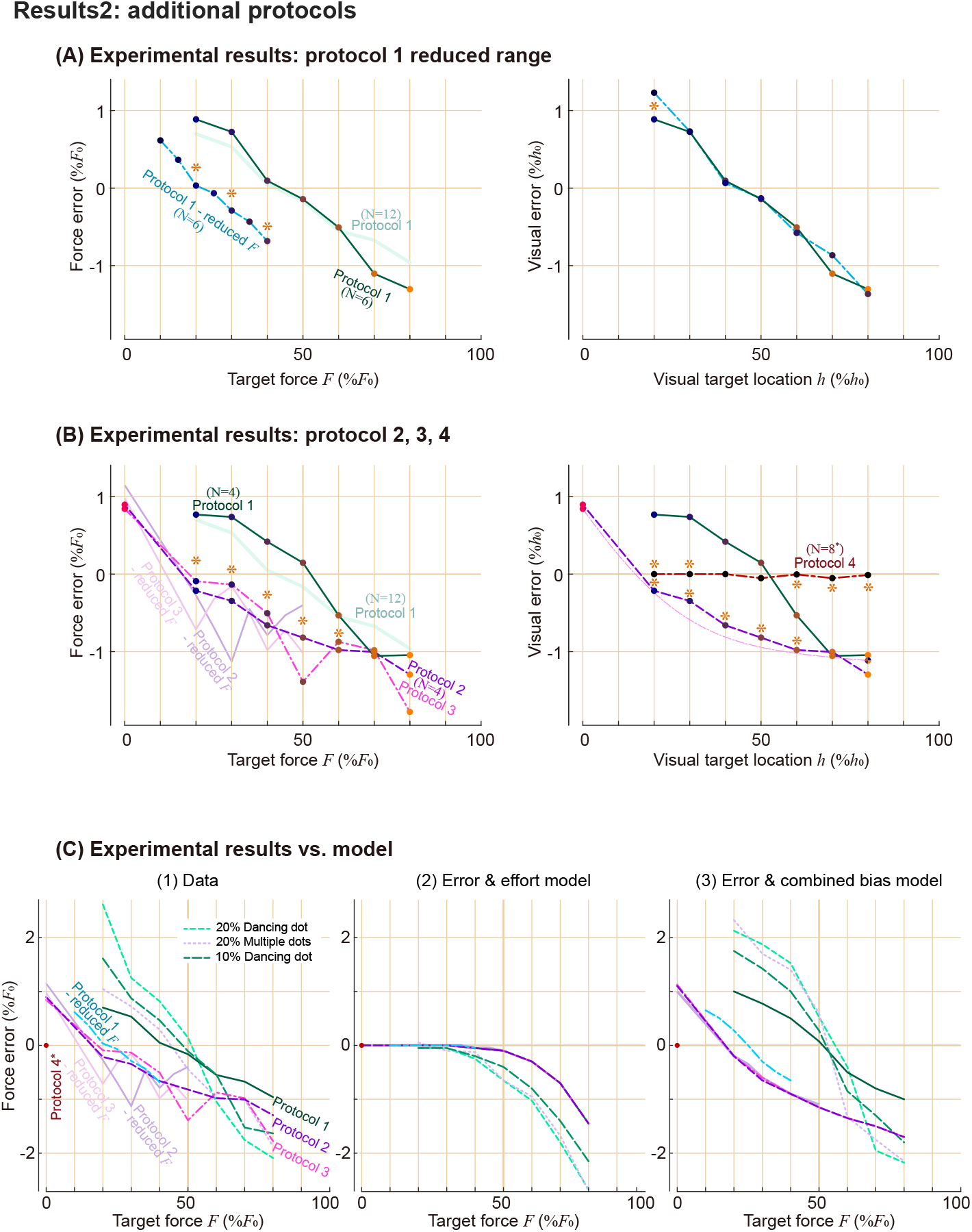
Experimental results of additional protocols and its comparison to model predictions. Different lines indicate the results of each protocol. As subsets of Protocol 1 subjects participated in additional tests, we represented the results of matching subjects in Protocol 1 as a thick green line here, while showing the results of all subjects as a dimmer line. Matching subjects result of Protocol 4 was omitted for simplicity, because the error was consistently very close to zero for everyone. (A) Protocol 1-reduced *F* trial had a distinct error trend compared to Protocol 1 in force domain, while its visual domain had a good match. Asterisks indicate statistically significant differences (p<0.05) between Protocol1-reduced *F* and Protocol 1 for matching conditions. (B) Protocols 2, 3, and 4 results, in force and visual domain. Asterisks indicate statistically significant differences between protocols and Protocol 1. (C) Comparison of (1) experimental results and (2, 3) models. Error-and-effort model and error-and-bias model were poor and good predictors of data.

### Protocol 2 and 3 shows that testing order affects the error, and that visual representation of the target force is not a good descriptor of error

We tested the same target force range as Protocol 1, but added zero force reset periods between non-zero force targets (figure 5A) to test the order effect. The difference between Protocol 2 and Protocol 3 was the visual representation of the target. In Protocol 2, visual representation was the same as the Protocol 1, i.e., the target location was moved proportional to the required force. In Protocol 3, targets appeared between only two locations, the lower one indicating 0 force and the higher one indicating various non-zero target forces. In this case, the ratio between force and target representation changed each time a new target was presented. On a different day from Protocol 1 testing, 4 participants came back to participate in Protocol 2 and Protocol 3.

## Results

Testing results suggest that visual representation of the target is also not a good predictor of the force tracking error. Despite different visual target representations, there were not significant differences between Protocol 2 and Protocol 3 in force-error relationship on all target force conditions (figure 6B). Those force-error relationships were significantly different from that of Protocol 1 in most of the target force range. The observations were similar in visual target location–error relationships. Error trends of Protocol 2 and Protocol 3 were similar to each other, but were different from that of Protocol 1. We also tested Protocol 2 - reduced *F* and Protocol 3-reduced *F* conditions, where we tested a smaller range of the forces, while not changing the mapping between force and its visual representations. There were no notable differences between Protocols 2 and 3 and their reduced *F* versions.

### Protocol 4 shows that visual bias does not explain the force bias

We asked subjects to track the target using a computer mouse, so that we could investigate the errors coming from visual aspects of the task. They did not need to keep exerting force to keep the mouse position, which is different from all other protocols where they needed to keep applying the force until a new target was shown. Eight out of twelve subjects performed this Protocol 4 experiment on their personal computers.

## Results

Participants performed the tracking task almost perfectly. Visual tracking errors were overall quite small, and the error relationship was again significantly different from that of Protocol 1 for most of the ranges (figure 6B).

### Model with force error and biases captured the experimental results qualitatively

We reject hypotheses that force or visual representation of the force predicts force tracking error, as force tracking errors were different for some protocols even for the same force or same visual target location. Any objective function model that mainly relies on mere force (e.g., figure 6C2) or visual representation to describe error trend would not be suitable to describe the observation. Error-and-bias model was better supported by the data. Error-and-central tendency bias model that fits Protocol 1 predicts the results of additional protocols fairly well (figure 6C3). The objective functions that minimized mean RMS error between mean data and mean model prediction on each force condition of different protocols were as follows (figure 7):

- Error-and-central tendency bias model:

**Figure 7.**
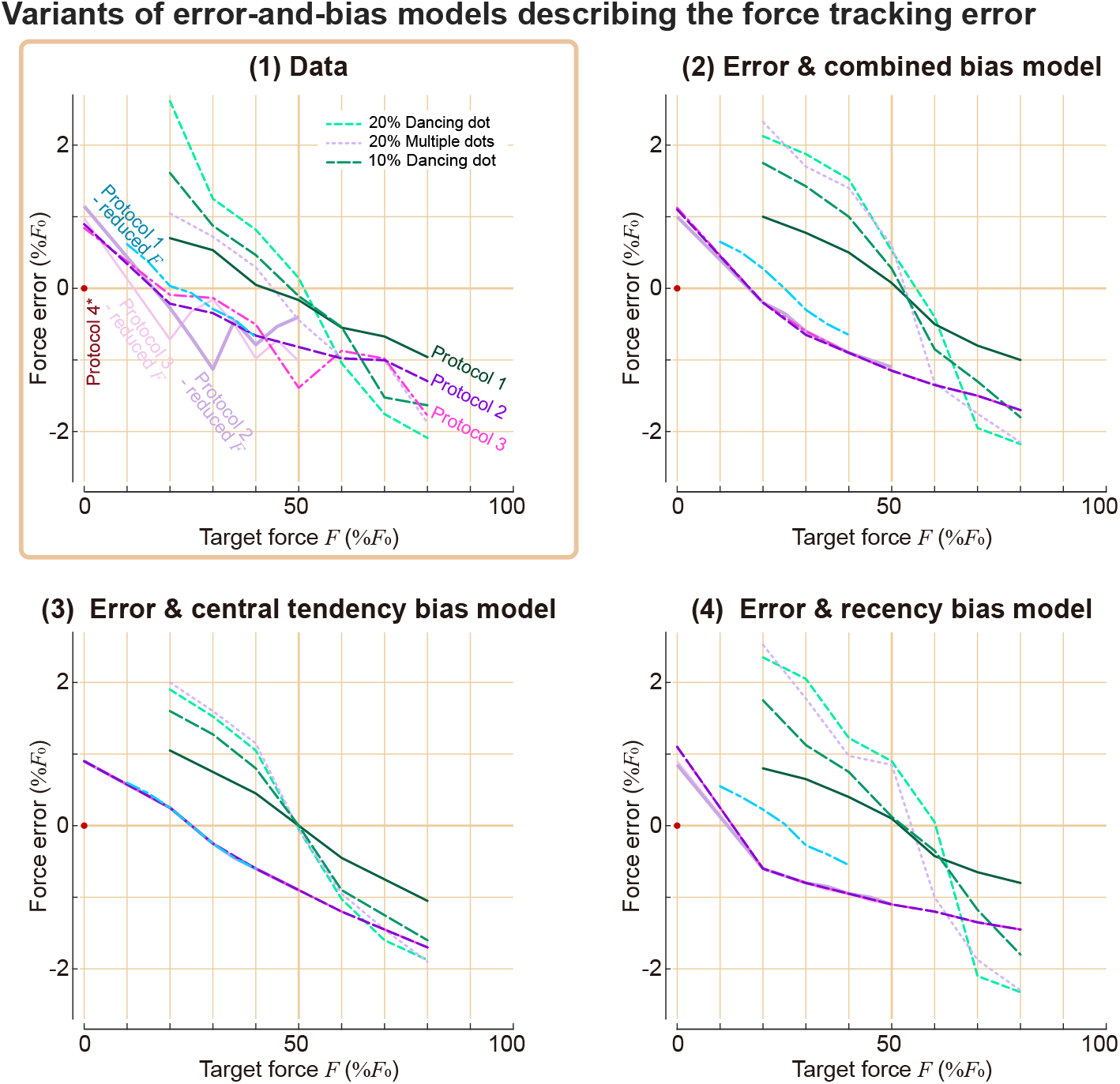
**Variants of error-and-bias models describing the force tracking error from various protocols.** (1) Experimental data from various protocols. Different lines indicate the median results of each protocol. (2)-(4) Models that best fit the overall experimental data, whose objective functions are weighted sums of the error terms and (2) both central tendency bias and recency bias terms, (3) central tendency bias term, and (4) recency bias term.

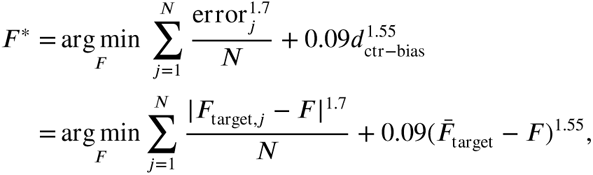

- Error-and-recency bias model:

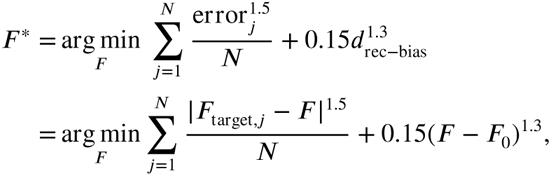

- Error-and-combined bias model:

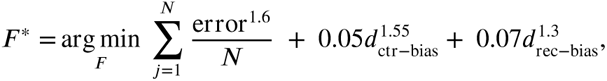

where *F* ^∗^ is the prediction of the applied force based on the given objective function and *F* is the applied force that is to be optimized. *F*_target,*J*_ is the force indicated by *J*-th dot in the sub-trial, when *N* is the number of dots presented. Bias-related terms *d*_ctr−bias_ is a distance to the “center” of the targets associated with central tendency bias, *d*_rec−bias_ is a distance to the recent action associated with recency bias. They are determined by 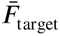, which is the average (or expected) target force of the whole trial consisting of many sub-trials, and *F*_0_, which is the force at the beginning of the sub-trial.

Details of the inverse optimization analysis are described in the Methods section.

### Changes in target forces describe error trends from all protocols

We further investigated how force tracking error is explained by a tendency to shift towards some point. As we pointed out earlier, the error-target force relationship (figure 8A) and the visual error-visual target location relationship had distinct patterns for each of the protocols. However, the expected change in target force (figure 8B) and change in target force from the recent past target (figure 8C) better describe errors across different protocols. These support the idea of people’s tendency to shift towards either the recent past action or the center of the task. Error-and-effort model could not explain the positive error, but error-and-bias models could explain it: central tendency bias predicts positive errors on lower-range target forces as the center of the task is higher than the current force, and recency bias predicts positive error on lower range target forces as their recent past targets were on average higher than the current target.

**Figure 8.**
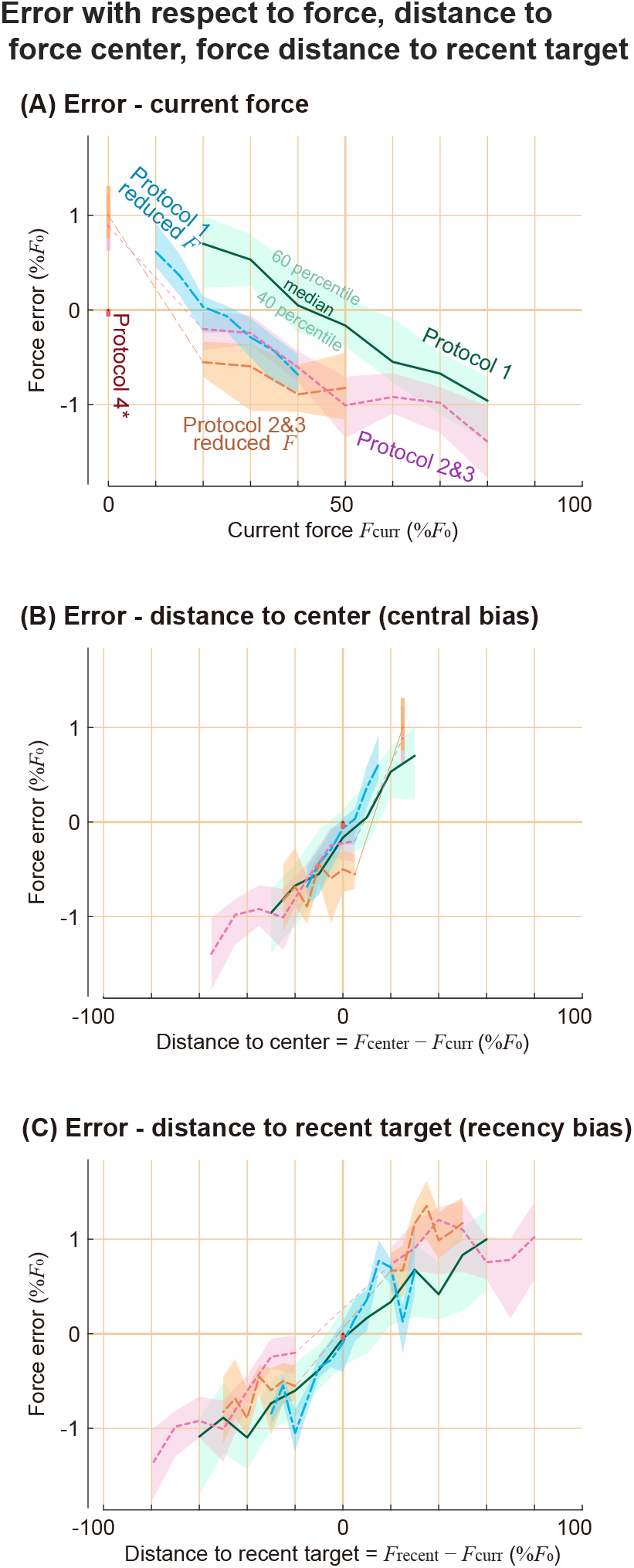
Error trends with force, force transition from the recent target, and expected future force transition. Force tracking error is plotted against (A) the current target force, (B) distance to the target center from the current target force, which is the expected transition, (C) distance to the recent past target force from the current target force, which is the transition from the past. Shaded areas represent 40 to 60 percentiles of the data at each condition, and the thick lines represent the median of the data. We showed visual error of Protocol 4 on the y-axis as a reference, because force error is undefined. The y-axis is for force error for all other conditions.

We defined recency bias as a function of *d*_rec−bias_, which is the distance to the recent past action, and central tendency bias as a function of *d*_ctr−bias_, which is the distance to the “center” of the targets or the average of the targets. From their definitions, the average of the *d*_rec−bias_ is close to *d*_ctr−bias_ except for the small discrepancies due to the order effect in Protocols 2 and 3. Therefore, models using recency bias, central tendency bias, or a combination of recency bias and central tendency bias, all yield similar overall error trends (figure 7C). We analyzed sub-grouped data to see their separate effects, and found that they both seem to exist, which is presented in the following section.

### Comparing central tendency bias and recency bias

#### Changes in target force affect error when the current target force is fixed

We grouped data of Protocol 1 – fixed dot target condition to investigate the recency bias effect. If recency bias does not have a notable effect, sub-trials that have the same target force will generally have the same errors, regardless of their recent past target. However, we do see a dependency on where the force started from, even if the target force is the same (figure 9A). Errors had negative correlations with force change, even when they were compared against sub-trials that had the same target force.

**Figure 9.**
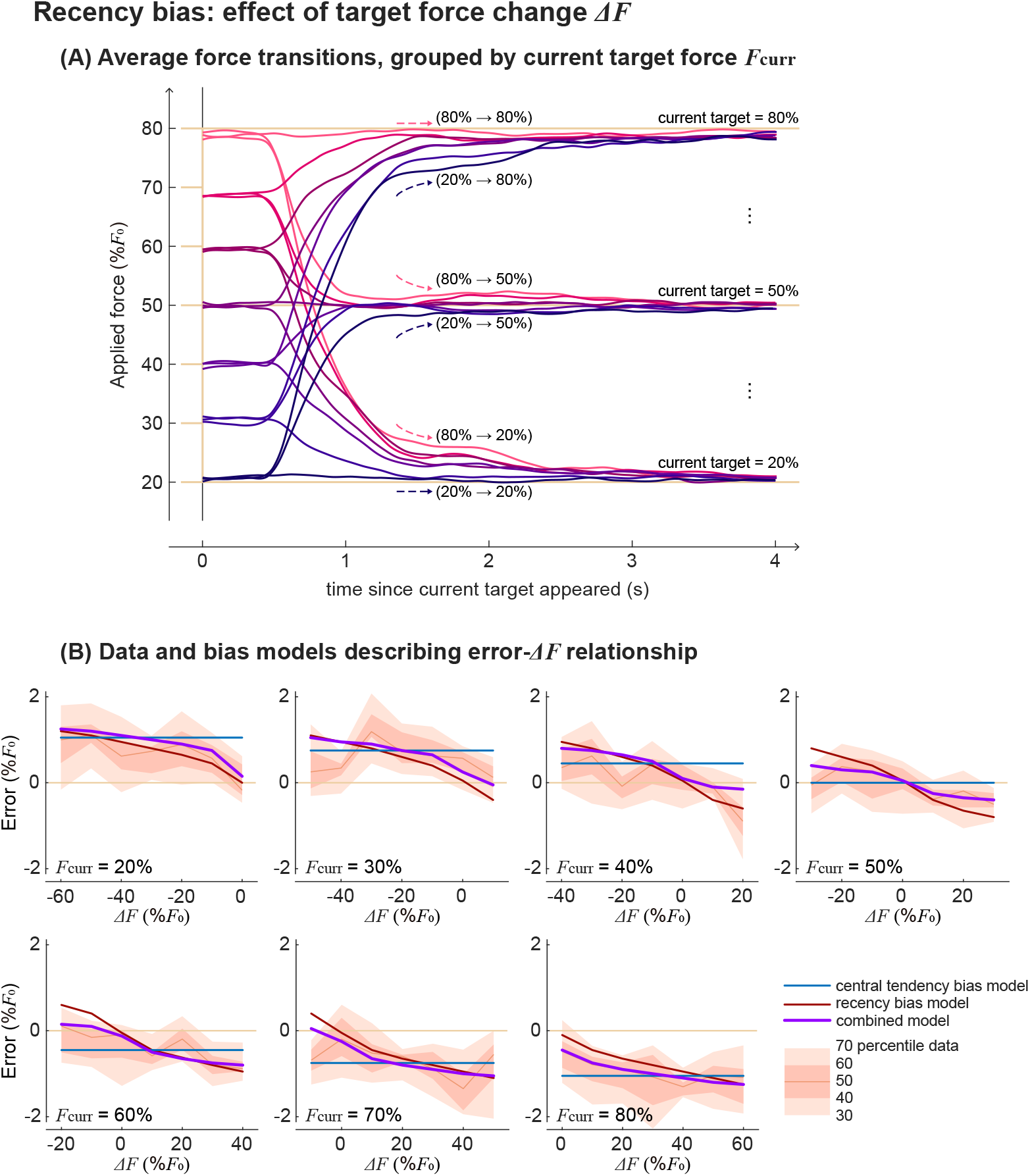
Protocol 1 sub-trials grouped into same target forces. (A) Averaged force time series grouped based on its recent past target and current target forces. Brighter and more red colors are for higher previous target forces, and darker and more blue colors are for lower previous target forces. (B) Force error as a function of target force change, among sub-trials that had the same current target force. Each sub-panel is for different current target forces. The shaded area represents 30, 40, 50 (=median), 60, 70 percentile data range. Blue, red, and purple lines are predictions using error-and-bias models with only central tendency bias, only recency bias, or central bias and recency bias combined.

We considered three types of bias models in the objective function: central tendency bias, recency bias, and a combined model that has both biases. While all three of them do similarly well at capturing the overall error trend with force, the central tendency bias model does not capture the negative correlation between force change and error when the target force is fixed (figure 9B). Recency bias and combined model are more suitable to describe this observation.

#### Current target force affects the error among sub-trials that underwent the same change in target force

We now grouped data of Protocol 1 – fixed dot target condition to investigate the central tendency bias effect. Here, we grouped sub-trials into the same target force changes, because analysis on the sub-trials that had the same target forces showed that change in target force affects the force error. If recency bias alone could explain the observations and central tendency bias has a negligible effect, groups of data that underwent the same target force change would have a similar error regardless of the current force requirement. However, we do see a dependency on the current target force level, when we compare sub-trials against other sub-trials that had the same target force change (figure 10A). Errors had negative correlations with target force in each of the groups that had the same change in target forces. We explained this observation in terms of central tendency bias, that people make positive errors when the center of the task is higher than the current target force, and make negative errors when the center is lower than the current target force.

**Figure 10.**
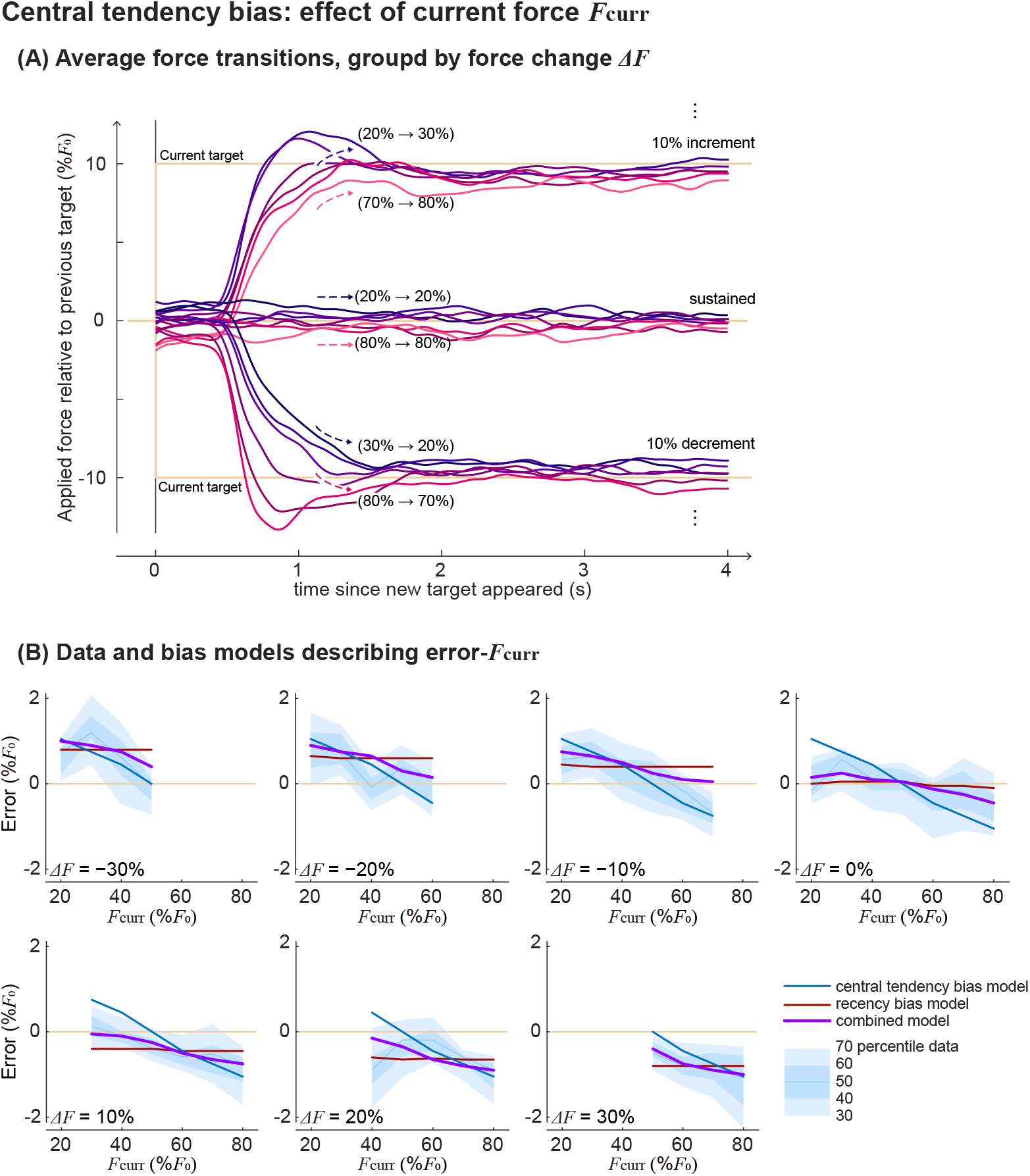
Protocol 1 sub-trials grouped into same changes in target forces. (A) Averaged force change time series, where the y-axis is a force difference with respect to the recent past target. Shown here are sub-trials that had 10%*F*_0_ increment, same, or 10%*F*_0_ decrement from the recent past target. Brighter and more red colors are for higher current target force, and darker and more blue colors are for lower current target force. (B) Force error as a function of the current force, among sub-trials that had the same change in the target force. Each sub-panel is for different changes in the target force. The shaded area represents 30, 40, 50 (=median), 60, 70 percentile data range. Blue, red, and purple lines are predictions using error-and-bias models with only central tendency bias, only recency bias, or central bias and recency bias combined.

We again considered three types of bias models in the objective function: central tendency bias, recency bias, and a combined model that has both biases. Since the change of target force is a variable that inherently co-varies with target force on average, all three models capture the overall error trend. However, the recency bias model does not capture the negative correlation between target force and error when the target force change is the same (figure 10B). Central tendency bias and combined model are more suitable to describe this observation.

### Purely visual tasks did not have the biases

We have shown that force or visual target locations alone were not good descriptors of error, but force bias could be a good descriptor. Then, if an error could be explained by force bias, there is a chance that it could also be explained by a visual bias. We therefore investigated whether visual bias could be an alternative explanation of the error trend we observed. Since the visual location of the target was proportional to the force in Protocol 1, visual bias could explain the error trend (figure 11 left) as much as the force bias does. The most striking counterexample is from Protocol 4, where subjects used a computer mouse to track the targets and did not need to exert force to maintain the pointing location. Although recency bias is commonly studied in visual tasks, we do not observe clear visual biases from Protocol 4 (figure 11 right). Subjects made almost no error for single fixed targets when they used a computer mouse to track the target. When they tracked a vague target which is a set of multiple dots, overall variability increased, but there was no significant trend in visual error, and its mean is not significantly different from 0 regardless of the visual target location.

**Figure 11.**
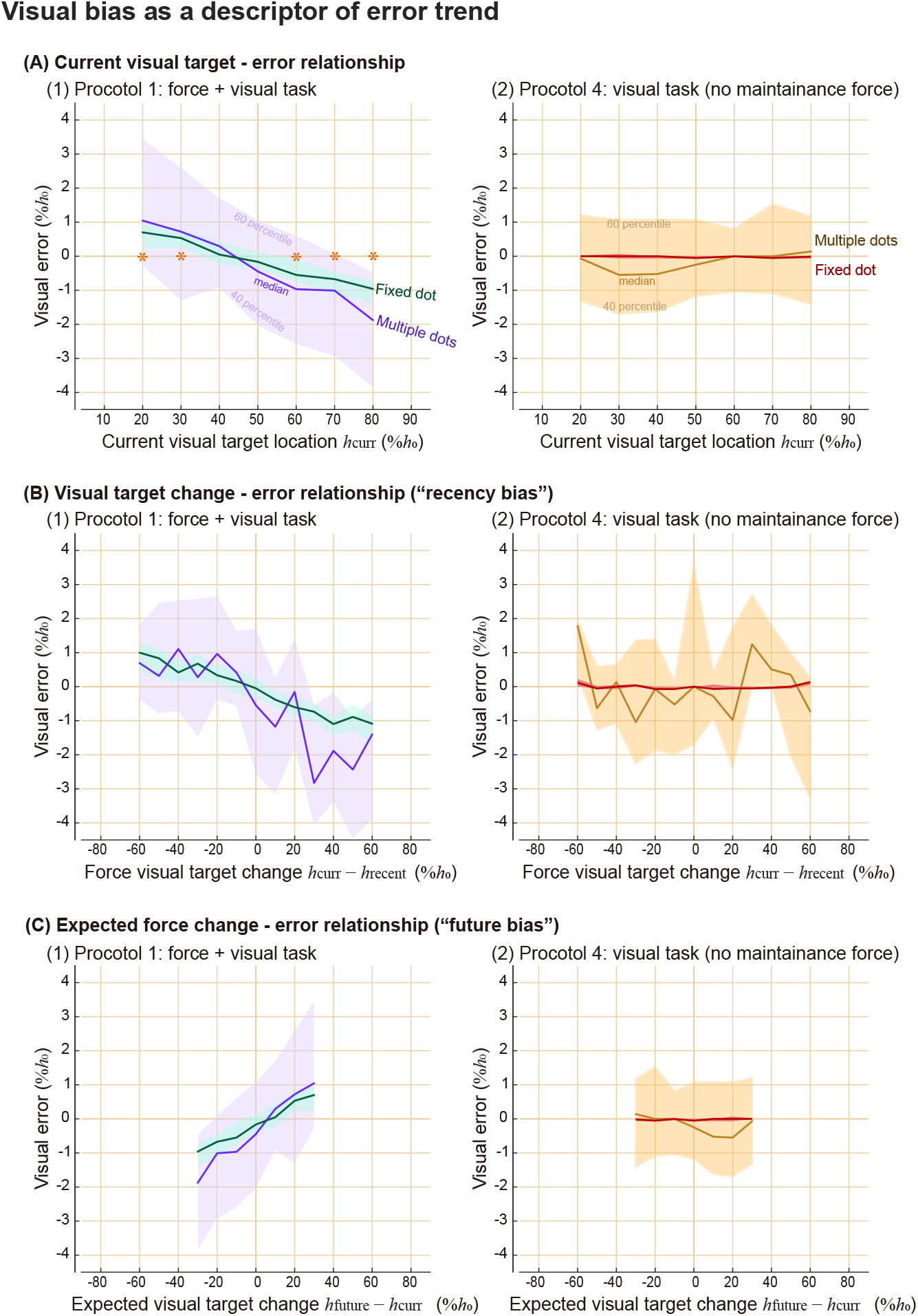
Tracking errors in visual domain, comparing Protocol 1 and Protocol 4. Results from target type of single dot and multiple dots are shown here. Shaded areas represent 40 and 60 percentile data, and thicker lines in the middle of the area are the median of the data. We represent visual error with respect to (A) the current visual target location, (B) the distance to the center of the visual target, (C) the distance to the recent past target location. Asterisks in the top row sub-panel indicate statistically significant differences from zero for both target types separately. There were no significant differences from zero from Protocol 4.

## Discussion

We measured force tracking errors while subjects performed tracking tasks of various protocols and modeled their behaviors using an objective function with an error term and force bias term. Our original aim was to measure the trade-off between task performance and effort, but we needed to include bias term(s) rather than an effort term in the objective function to explain the result. The main discrepancy between the observation and our original expectation was that people made consistent positive force errors in some ranges, which is not explainable by either effort-saving tendency or error-reducing tendency. Central tendency bias and recency bias, which means the tendency to shift towards the center of the tasks and towards the recent action, seem to explain the data well when combined with an error model. To our knowledge, these biases in force tracking tasks have not previously been observed and modeled.

In certain modeling scenarios, refining the control objective function, as demonstrated in this study, can yield significant benefits, while in others, the advantages may be less evident. The usefulness of such refinement may depend on the level of detail at which one examines the phenomenon, and on practical considerations such as computational requirements and analytical solvability. On the other hand, understanding the human tendency to make biases while performing motor tasks could have broader applications in various fields, including human-machine interaction and human factors engineering. Additionally, exploring the connection between these biases and neuromuscular disorders could also provide deeper insights into motor control and hold the potential for various clinical applications, including diagnosis and rehabilitation.

We initially attempted to model the force tracking error using an effort model because it is a common expectation that motor control is usually done in an energetically optimal way. The observation that people apply more force than needed in the low-medium force range seems to be an example that people did not optimize for energy, at least during the experiment. However, could it turn out to be energetically cheaper to make such positive biases? For example, if there is a huge cost associated with force changes, it could be more efficient to spend slightly more energy on maintaining force and save a bigger energy by reducing a force change. Since we did not directly measure the energetic cost during the experiment, we cannot conclusively claim that people’s behavior was energetically suboptimal. However, our speculation is that energy saving is not a primary explanation of the biases, because 1) the transient cost people save seems to be small compared to what they spend throughout the sub-trials while maintaining the force, 2) usually “fast change” is regarded energetically costly, but people did not seem to slow down the transient even when there was no explicit speed requirement, suggesting that transient cost is not of significant consideration, and 3) people often did reach a lower force than the target, and increased the force to again to make positive force error at the end (***Box 1***).

We modeled force tracking error as an outcome of competing goals of reducing error and biasing towards a center, but there could be alternative models to capture the same phenomenon. We used our bias models to predict errors from additional protocols, but the model is still descriptive in the sense that we did not test for the underlying mechanism or causality of such bias. Our description of biases suggests that people tend to reduce force change, either from the past by making an incomplete shift, or toward the future by making an anticipatory error towards the expected target. However, since we have not proven whether the reduction in force change is the primary cause of the errors, it may turn out to be a secondary change following another one. It would be interesting for future work to investigate the causality of the biases.

Including more terms in the objective function model could capture a broader range of phenomena, but it could also make the inverse optimization problem more underdetermined, potentially leading to an unreliable solution. Central tendency bias and recency bias exhibit similar effects on the overall trend predictions in our experimental designs; thus, one may not need to model both biases to explain some behaviors, depending on their goals. We did not particularly design experiments to distinguish these two biases, and there were multiple sets of solutions yielding similarly good inverse optimization results (figure 7). Our subgroup analysis suggests that both seem to have an effect, but we do not claim that we quantitatively verified recency bias and central tendency bias. It would be interesting to design experiments to further disambiguate the biases that originate from past actions and originate from anticipatory actions for the future. For example, one could design a trial order that cycles through three stages, two random non-zero target forces followed by one zero target, so that the average past action is distinct from the average future actions. Even the formulation of the error-reducing tendency must have been an oversimplification if the aim was to closely understand what people perceive as a target. Error function could have a different form depending on the number of dots presented as a target, or could take into account how individual subjects subjectively defined the tasks.

There could be various biases in human motor control, and some of them could happen immediately without task-specific experiences while some of them may need learning of a task. Some of the biases could take a long time to change their properties, while others may rapidly adapt to each of the tasks. We assumed that subjects readily had an expectation of the center of the target when we modeled central tendency bias, but further research is needed to justify this in depth. We chose to model the central tendency bias this way because we had speculated that the bias does not seem to rely on learning from each trial, because force tracking errors seemed to happen instantly when testing began, and because we do not observe clear changes over the course of the trial. However, we do not have strong evidence to show whether subjects make biases based on their experience of the trial or expectation of it, or based on the combination of them or even neither of them. It is conceivable that people may start making biases based on their prior expectations and refine the expectations as they have more experience with the task. It would be interesting to design experiments that give subjects a more different impression of the task from its actual composition, and to see which one is a better predictor of the behaviors.

One of the reasons that biases in force tracking tasks were not commonly recognized in biomechanics studies might be due to the specifics of the common experimental designs. In biomechanics literature, it is common to 1) measure force transients that start from rest, and 2) report the absolute value of the error, which omits the directionality of the error. Researchers sometimes describe that people are less accurate at producing bigger forces. Our findings suggest that this description might be highly context-dependent, and there could be a more general description of the phenomenon. Our error-and-bias model predicts that absolute error is minimized near the center of the tasks (figure 12A), rather than monotonically increasing with the force magnitude. Despite that overall root-mean-squared (RMS) errors in experimental results were shifted up compared to the model predictions (potentially due to human variabilities), our model captures the overall trend of error (figure 12B) that had minima near the center of the task.

**Figure 12.**
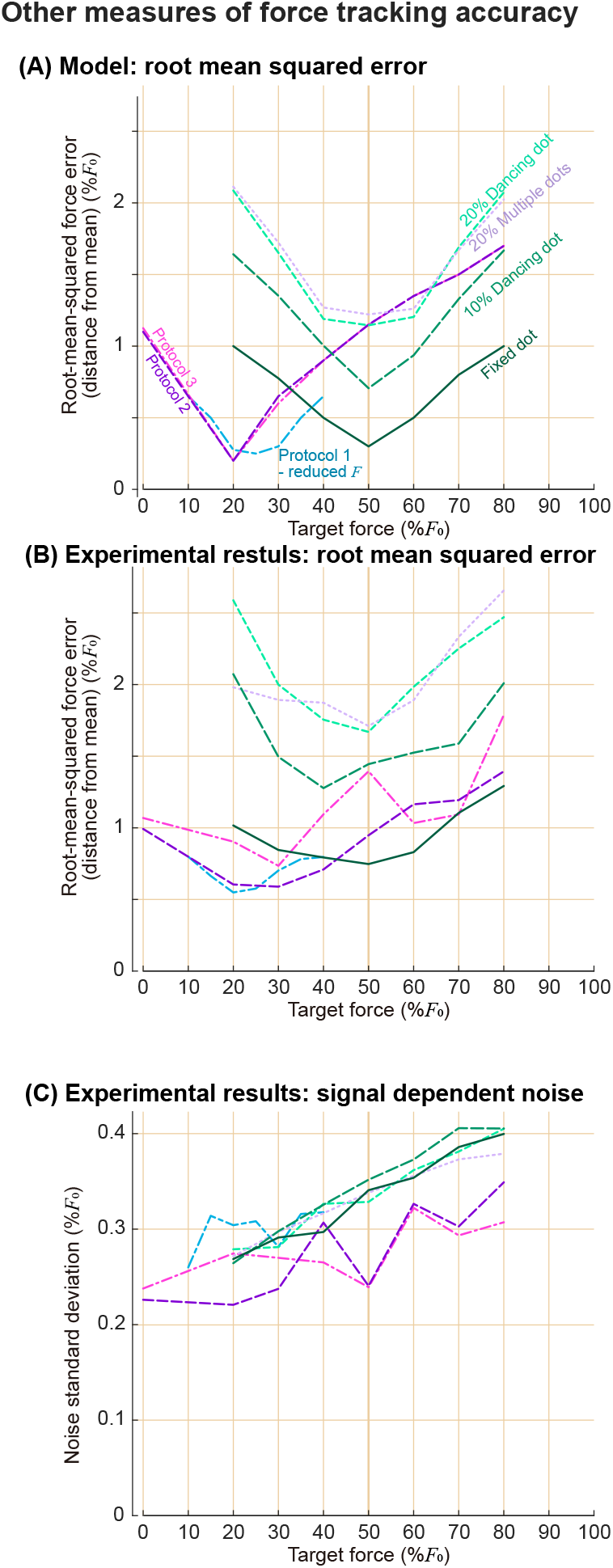
Root-mean-squared error and signal-dependent noise during force tracking tasks. For root-mean-squared (RMS) error, we use error defined as a distance between the mean of the target distributions and the average force applied during the last 0.5 seconds of the sub-trials. We show RMS error that (A) model predicted and (B) measured from experiments. Lines of different colors indicate different protocols and target types. The target force on the x-axis indicates the mode of the target distributions. (C) The standard deviation of the higher frequency component of the force, with respect to the target force. This higher frequency component of force is referred to as motor noise in some literature, and its dependency on the target force is an example of signal-dependent noise. Lines of different colors indicate different protocols and target types.

### Box 1

**Box 0—figure 1.**
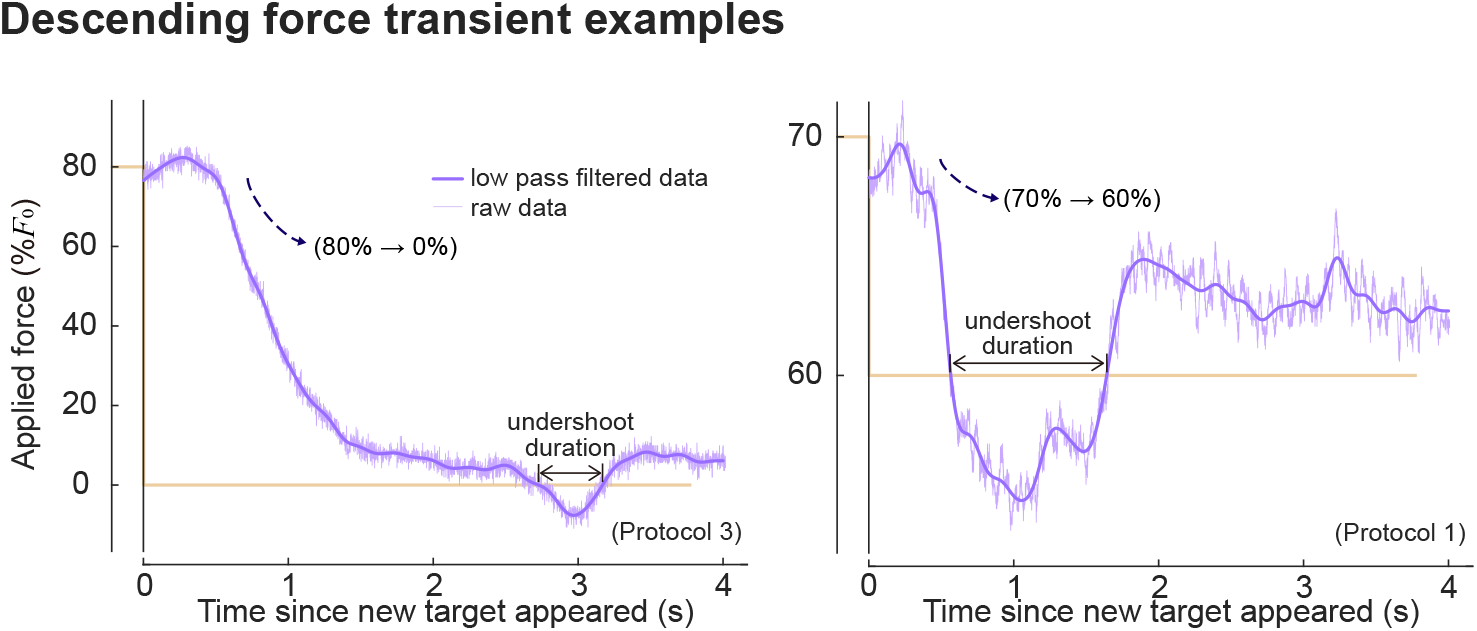
Example force time series during sub-trials that had descending target forces. Dim horizontal lines indicate the recent past target and the current target. Lighter purple lines show raw data, and darker purple lines show low pass-filtered data that was used in undershoot analysis. We defined undershoot duration as the duration where the applied force was lower than the target force during descending sub-trials.

Undershoots in descending sub-trials (where the current target force was lower than the recent past target) are particularly interesting because they provide insights into the underlying reason for the positive force errors.

Undershoots in descending sub-trials (where the current target force was lower than the recent past target) are particularly interesting because they provide insights into the underlying reason for the positive force errors. Overshoots and undershoots that go past the target were quite common when we looked at individual sub-trials, although they were less obvious in averaged force transient responses (figure 9 and figure 10) due to the large variabilities in both amplitude and timing of the responses between sub-trials. In Protocols 1, 2, and 3 full range and reduced range trials, 72-90% of descending force transients had undershoot of more than 0.2 seconds. Among all descending transients, 67-85% of sub-trials had positive tracking errors at the end. Among descending transients that ended up having positive errors, 68-86% had undershoot of more than 0.2 seconds. There were many sub-trials where the force recovered back to a higher force after an undershoot during descending force transients like the sub-trials shown here, where it did not seem like an unavoidable by-product of a fast response due to physiological constraints, nor a strategy to save transient energy. This suggests that positive biases on low-medium range target forces are not for energetic advantages but for some other benefits.

Our results are still consistent with the previous studies, when we look at the non-zero sub-trials of our Protocols 2 and 3, which were always followed by zero force sub-trials. RMS errors indeed roughly increased with the force magnitude in these cases. Directionality of the error during these sub-trials was also consistent with previous studies: we predict and measure negative errors, and previous studies also usually measured negative force tracking errors (e.g., directly reported in ***Todorov (2002)***, could be inferred from figures in (***Kudzia et al., 2022***)). However, what had been usually not measured and reported in previous studies is that RMS force error during zero force sub-trials increases again (figure 12A), and that people tend to make a positive error for this range. The RMS error trend was even more dramatically different from the monotonic increase when we consider Protocol 1 results, where target forces were not reset to zero force.

This leaves us a question of whether we could truly eliminate order effects in scientific measurements. As we pointed out, even separating trials by resting periods and having a “fresh start” could give us a wrong impression of the phenomenon, because starting from zero is one of the special cases as well. It would be important to be 1) consistent and clear with what we measure, 2) recognize that what we observe might be a special case of a more general phenomenon, and 3) try to distinguish co-varying variables if possible. For example, force magnitude and force increment co-varies if tests always begin from zero, but they did not co-vary in our Protocol 1, and provided us with a more general view of the nature of force tracking errors. On the other hand, signal-dependent noise, which is defined as the standard deviation of force after removing a low-frequency component, had a closer match between trials and seemed to increase with force (figure 12C; consistent with ***Jones et al. (2002)***). This quantity seems to be less context-dependent than force error, and thus, has a higher chance of being explained by simpler physiological properties such as muscle characteristics or nervous system dynamics. However, what we measured could still be a partial description of a more general phenomenon, as it always is in science.

It was interesting to us that we did not observe a significant bias when people performed an almost fully visual task (figure 11), considering that many studies on recency bias and central tendency bias are measured from visual pattern recognition tasks. It is conceivable that human minds would have various biases without involving the muscular system. On the other hand, some biases may be caused by more passive and physical properties of the body without involving perceptions and judgments, and some biases might be caused by combinations of them. From studying the biases of force tracking tasks, we became curious if there is a chance that some of the perceptual biases could be attributed to motor control biases more than researchers commonly thought. For example, ocular motor control may have a similar bias as what we observed in this study that involved arm muscles, and that could result in a bias in visual sensation. There could be biases caused during measurement procedures as well, because subjects were often asked to indicate their perception of patterns using some physical device by manipulating it, which involves some degree of motor control.

Force bias does not seem to be a commonly recognized phenomenon in an isolated motor control setting, but we could relate it to a broader field of motor control. In a singing study, it was reported that people tend to compress the pitch shifts, and “bad singers” tend to make more such compression (***Pfordresher and Brown, 2007***). In speech studies, there is a window model suggesting that people make minimal shifts between acceptable windows of some variables (***Keating et al., 1990***). Moreover, when we consider the recency bias in broader terms that people tend to keep acting similarly, there are more such examples in biomechanics studies. For example, walk-to-run and run-to-walk speeds are reported to be different (“hysteresis effect in speed for the walk-run transition”, ***Diedrich and Warren Jr (1995)***), which is not simply explained by the idea that animals choose the most economic gait at a given speed. People change their behavioral mode between one-handed grasping and two-handed grasping depending on the size of the object, and it is reported that there is an order effect in such shifts (***Frank et al., 2009***). Some researchers attribute these biases to the “economy”, that people save energy by reducing transitions, although this claim is usually difficult to be directly tested. We hope that an extension of our current study could give insights into these broader fields of behavioral studies.

To conclude, we examined which objective function best describes human force tracking errors. In many motor control studies, objective functions are modeled to include error and effort terms, typically in quadratic forms. However, we designed an experiment to test them and indeed found a different formulation. The exponent on the error we found was smaller than 2, and bias term(s) were needed instead of an effort term in order to capture the behavior that was consistent across participants. Our findings on biases suggest that biological motor control, even in a simple isometric force production task, can be highly context-dependent, and commonly hypothesized formulation might not be adequate to represent the phenomenon. As we continue to explore the intricacies of human force control, further research may offer valuable insights into our understanding of human movement behaviors and underlying principles of the nervous system.

## Methods

### Experimental procedure

Subjects (*N*_subjects_ = 12; 4 Female, 8 Male) participated in Protocol 1, and a subset of them participated in additional studies later. This human subject research was reviewed and approved by The Ohio State University’s Institutional Review Board, and all subjects participated with written informed consent.

We measured elbow height from the ground and subtracted half of the subject’s forearm length to set a platform height they pressed onto, which made the forearm angle about 30 degrees below the horizontal line. After setting the force platform, we measured the individualized force target range *F*_0_ by asking the participants to apply a force that they could comfortably hold for 30 seconds. We encouraged participants to apply reasonably high force, and reduce it until it felt comfortable to hold. Real-time force feedback was given on the monitor, and we encouraged them to control the feedback bar location. After measuring the force range, we verified again that subjects could comfortably exert the force for 30 seconds. The process was repeated until we found the force that the subject was comfortable with producing for an extended amount of time.

After giving participants a verbal description of the tasks, we provided them with practice trials, which were shorter versions of the trials so that they could see the test procedure and try the tasks. We introduced different target types in turn – fixed single dot, then single dancing dot, and then multiple dots. The target changed color between blue and red in each sub-trial, and all participants confirmed that the colors were significantly different for them. We instructed the participants that the target location changed every 4 seconds with the change of the target color, and asked them to apply force so that the force feedback bar points to the target. For vague targets, we told them that the dots are dancing around some fixed point and are not moving away as long as the color stays the same, and asked them to point where they perceive as a target from the overall impression of the dots. We explicitly asked them not to chase individual occurrences of the dots, but to perceive its center and place the bar there. Participants practiced each of the target types for at least 5 sub-trials, and we repeated the practice if subjects desired to do so or were unclear about the task.

Each trial was composed of 74 sub-trials. The first three trials were excluded from the analysis to make sure that subjects adapted to a different target type. The following 70 conditions covered all possible combinations of force level and distribution parameters in a random order, which is described in the following paragraph. After these 70 sub-trials that are used for analysis, the last sub-trial of each trial was 100%*F*_0_ one, from which we checked that subjects were still capable of producing more than 80% of *F*_0_. 4 seconds of 74 sub-trials took about 5 minutes.

We tested 20, 30, …, 80% of *F*_0_ in Protocol 1, and used distribution parameter *α* = −2, −1, −0.5, −0.1, −0, +0, +0.1, +0.5, +1, +2 to generate vague targets, resulting in 7 force levels × 10 distribution parameters = 70 combinations. For the fixed single dot condition, we measured each force level 10 times to ensure the same number of sub-trials. In Protocol 1 - reduced *F*, force levels we tested were 10, 15, …, 40 %*F*_0_. In Protocols 2 and 3, non-zero forces were 20, 30, …, 80%*F*_0_, and each force level was measured 5 times, again resulting in a total of 70 sub-trails including zero force periods. In Protocol 2 and 3 - reduced *F* conditions, non-zero force levels were 20, 25, 30, …, 50 %*F*_0_. Protocol 3 had binary visual target locations, which were the same as the target for 0 force and 80%*F*_0_ in Protocol 2.

We defined the distribution parameter *α* as follows. Positive non-zero distribution parameters correspond to a shape parameter of Pareto Distribution truncated between 0 and 5 (***Zaninetti and Ferraro, 2008***). The range was scaled to match 10% and 20%*F*_0_. We generated distributions of negative non-zero *α* same as a positive *α*, except that the whole distribution was flipped so that the tail of the distribution goes the opposite way. Two symmetric distributions were also used: truncated uniform distribution was referred to as *α* = −0, and truncated normal distribution that had similar standard deviation as other Pareto distributions were referred to as *α* = +0 in this study as special cases of skewed distributions.

Six of the subjects had force feedback that went down as they applied force, and six of the subjects had force feedback that went up as they applied force. We did so to see if there was a visual bias linked to a vertical directionality. We did not observe a notable difference between the subjects of each feedback directionality. Each time before the test started, we asked subjects to place their hands on the platform and relax their arm muscles, try not to exert active force onto the platform, and let the platform take the weight of the hands. This hand weight was set to zero force so that participants did not need to actively spend energy lifting up their hands to match low force requirements.

Protocols 1 and 4 involved testing vague targets. In Protocol 1 – dancing dot target condition, a dot appeared within 10% and 20%*F*_0_ range, and in Protocol 1 and 4 – multiple dots conditions, 240 dots appeared within 20%*F*_0_. We tested all target types in a random order two times. Other protocols were measured once per protocol. Subjects took about 2 minutes of rest between trials and occasionally took longer breaks as desired.

### Data collection and processing

Custom MATLAB software was used to show targets and force feedback on the screen. A force plate (Bertec Corporation, Ohio, USA) mounted on the ground was used to collect force data. Force plate data was collected through a motion capture interface (Nexus, Vicon, Oxford, UK) that was relayed to MATLAB software. Target and force feedback were updated at 60 Hz, which matched the monitor refresh rate. Force was collected at 8 times faster speed, which is 960 Hz. Multiple dots changed their horizontal positions in every frame. The target radius was about 0.7%*F*_0_.

The mean value of force during the last 0.5 seconds of each sub-trial was defined as the steady-state force value, and this value minus the target force was defined as force error. We normalized force error by *F*_0_. The mean value of force during the first 0.1 seconds of each sub-trials was defined as an initial force, and was used to calculate recency bias. To study signal-dependent noise, we obtained a fast component of the force change by subtracting a slow frequency response from a faster frequency response. Fifth-order Butterworth low pass filters with a cut-off frequency of 25Hz and 5Hz were applied to force data using zero-phase filtering in order to calculate these slow and faster responses. Standard deviation of this fast component during the last 0.5 seconds was reported as a “noise” to investigate signal-dependent noise (figure 12).

### Protocol 4 - test using a computer mouse with negligible forces

Custom MATLAB software was used to conduct tests of Protocol 4. We kept the testing interface as similar as possible to other protocols, except that the vertical position of the horizontal bar that used to indicate the applied force was changed to indicate the vertical position of the mouse pointer on the screen. The cursor was hidden and subjects could only see the height of the horizontal bar as they moved the computer mouse.

We used a MATLAB callback function that responded to mouse position changes. Trials were similar to those of Protocol 1, in terms of test duration and conditions. We tested a fixed single dot and multiple dots using this interface. Unlike Protocol 1, multiple dots did not change their horizontal locations in Protocol 4, because we could not guarantee a constant refresh rate through the interface we employed. We recorded the time and position of the mouse pointer each time there was a change. Clicking was not required to perform the task.

### Predictions for additional protocols based on the bias terms

We considered two types of biases in this paper. Predictions of the force tracking error based on central tendency bias and recency bias are as follows.

#### Predictions based on central tendency bias

We modeled a tendency to shift towards the “center” of the tasks, where “center” was defined as the mean of the target forces throughout the trial. The mean of the tasks in Protocol 1 was 50%*F*_0_, and the mean of the tasks in Protocol 1-reduced *F* was 25%*F*_0_. Using this model, we expected that errors would in general cross 0 around these force levels in each of the protocols. In Protocol 2 and 3, since half of the subtrials were at 0%, the mean of the targets was 25%*F*_0_. We expected that zero force sub-trials would have positive errors, and non-zero force sub-trials would have more negative errors as the target force increased. We used the same center for Protocols 2 and 3 – reduced *F*, because subjects were unaware of the reduced force range.

#### Predictions based on recency bias

We modeled a tendency to shift towards the recent past action. In Protocol 1 and Protocol 1-reduced *F*, the prediction based on the recency bias model was that lower target forces would have positive errors, because their recent past force is higher than their current forces on average. Similarly, the prediction on higher forces was that they would have negative errors, and medium-range forces would have errors near zero. This distinction of low, medium, and high forces was all relative to the force range of each trial in this bias model. Similar to the central tendency bias model, we expect that errors would cross zero near 50% *F*_0_ and 25% *F*_0_ respectively for Protocol 1 - full range and reduced *F* conditions.

In Protocols 2 and 3 where there were zero reset periods, we expected that zero force subtrials would have positive errors because their recent past forces were always higher than the current force. Similarly, we expected that non-zero sub-trials would have more negative error as the target force increased, because their recent past forces were always zero. Protocols 2 and 3 – reduced *F* were expected to have similar error trends as their full *F* range versions. The only difference could be on zero force sub-trials, because their average distance to the recent past target was smaller in reduced range protocols than in full-range protocols, thus resulting in a slightly smaller positive error on zero force sub-trials.

Both central tendency bias and recency bias models predict similar overall error trends because their expected values are similar to each other. Both models predict that neither error-target force relationship, nor visual error-visual location relationship will be consistent across different protocols, because bias is dependent on force and the context of the experiments. In addition, both bias models predict that there is a bias in the force domain, not in the visual domain; thus, we predicted that Protocol 4 will not have such bias because the task does not require maintenance force.

### Inverse optimization

We did grid search to perform inverse optimization to select hyperparameters of objective function models. We used a similar mathematical formulation to represent both central tendency bias and recency bias, which was:

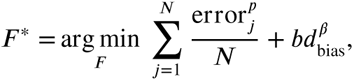

where *F* ^∗^ is the prediction of the force which minimizes the given objective function, error_j_ is absolute distance between *J*-th dot and the force applied, summed over number of target dots *N*. *d*_bias_ is absolute distance from the applied force to the bias center. It is the distance to the perceived average of the target forces for central tendency bias, and distance to the initial force at the beginning of the sub-trial for recency bias.

We searched for error exponents *p* between 1.4 and 1.9 range by the increments of 0.1, bias weightings *b* between 0.01 and 0.25 by the increments of 0.01, and bias exponent *β* between 1 and 2 by the increments of 0.05. For each combination of these hyperparameters, we calculated the model prediction of each sub-trials. The model predicts the forces that minimize the given objective function. To avoid local minima issue and to improve the computational efficiency, we evaluated the objective function within ±30%*F*_0_ around the target by the increments of 0.05%*F*_0_, and found the minimum value among them.

After calculating predictions for each sub-trial and each combination of hyperparameters, we selected the set of hyperparameters that produced a similar error-force relationship as the experimental results. Since error trend we aimed to model was in a relatively small magnitude compared to large inter-subject and intra-subject variabilities, we selected an objective function that captures the overall behavior. The objective functions that minimized mean RMS error between mean data and mean model prediction on each force condition for the entire protocols were:

- Error-and-central tendency bias model:

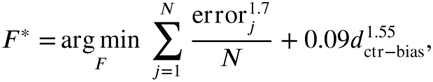

- Error-and-recency bias model:

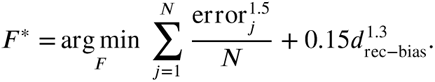

After performing inverse optimization on the central tendency bias model and recency bias model separately, we confirmed that they indeed have a similar formulation and have a similar effect of the force error predictions. We performed an inverse optimization on a combined bias model while keeping some of the hyperparameters fixed based on these results. We chose to fix some parameters to avoid overfitting, because we have confirmed that two biases have similar effects on the overall results, and human data is already very noisy.

We scanned the error exponent *p* between 1.4 and 1.9 by the increments of 0.1, while changing two bias weightings between 0.01 and 0.25 by the increments of 0.01. The bias exponent *β* was fixed to the optimal value that was found earlier. The objective function that minimized RMS distance between mean data and mean model predictions on error-force relationships was:

- Error-and-combined bias model:

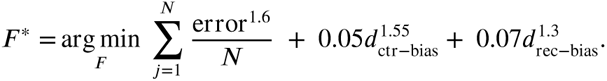

## Acknowledgments

HXR was supported by the U. Calgary Department of Biomedical Engineering. MS was supported in part by NIH-R01GM135923-01 and NSF SCH grant 2014506.

